# Ligand-specific endothelial Notch signaling coordinates brain pericyte specification and expansion

**DOI:** 10.64898/2026.10.05.756680

**Authors:** Huaxing Zi, Chi Dang, Weiyi Sima, Xiaoyu Li, Guozhu Ning, Li Sun, Qiang Wang, Jingjing Zhang

**Affiliations:** Affiliated Hospital of Guangdong Medical University & School of Medical Technology, Guangdong Medical University, Zhanjiang 524001, China; Dongguan Innovation Institute, Guangdong Medical University, Dongguan 523808, China

**Keywords:** brain pericytes, endothelial cells, Notch signaling, Jag2b, Dll4, Piezo1, zebrafish

## Abstract

Pericyte specification and expansion must be coordinated with vascular growth to establish a functional cerebrovascular network, yet how endothelial signals control these distinct developmental transitions remains incompletely understood. Here, using genetic manipulation, single-cell transcriptomic analysis, and longitudinal *in vivo* imaging in zebrafish, we identify ligand-specific endothelial Notch signaling as a mechanism coordinating brain pericyte development. Endothelial Jag2b is required for efficient pericyte specification from pericerebral progenitor pools and subsequently promotes Notch-dependent pericyte proliferation and population expansion. By contrast, endothelial Dll4 restrains pericyte proliferation and functionally opposes Jag2b-dependent expansion, demonstrating that distinct endothelial Notch ligands generate different developmental outputs. Endothelial Notch activation increases *jag2b* expression, and genetic interaction analyses establish Jag2b as an important mediator of Notch-dependent pericyte expansion. Moreover, genetic disruption of *piezo1* reduces brain pericyte abundance, whereas endothelial *jag2b* expression rescues this phenotype, placing Jag2b downstream of Piezo1-dependent hemodynamic signaling. Together, these findings reveal a ligand-specific endothelial Notch program that integrates hemodynamic cues with endothelial–pericyte communication to coordinate the specification and expansion of brain pericytes during cerebrovascular development.

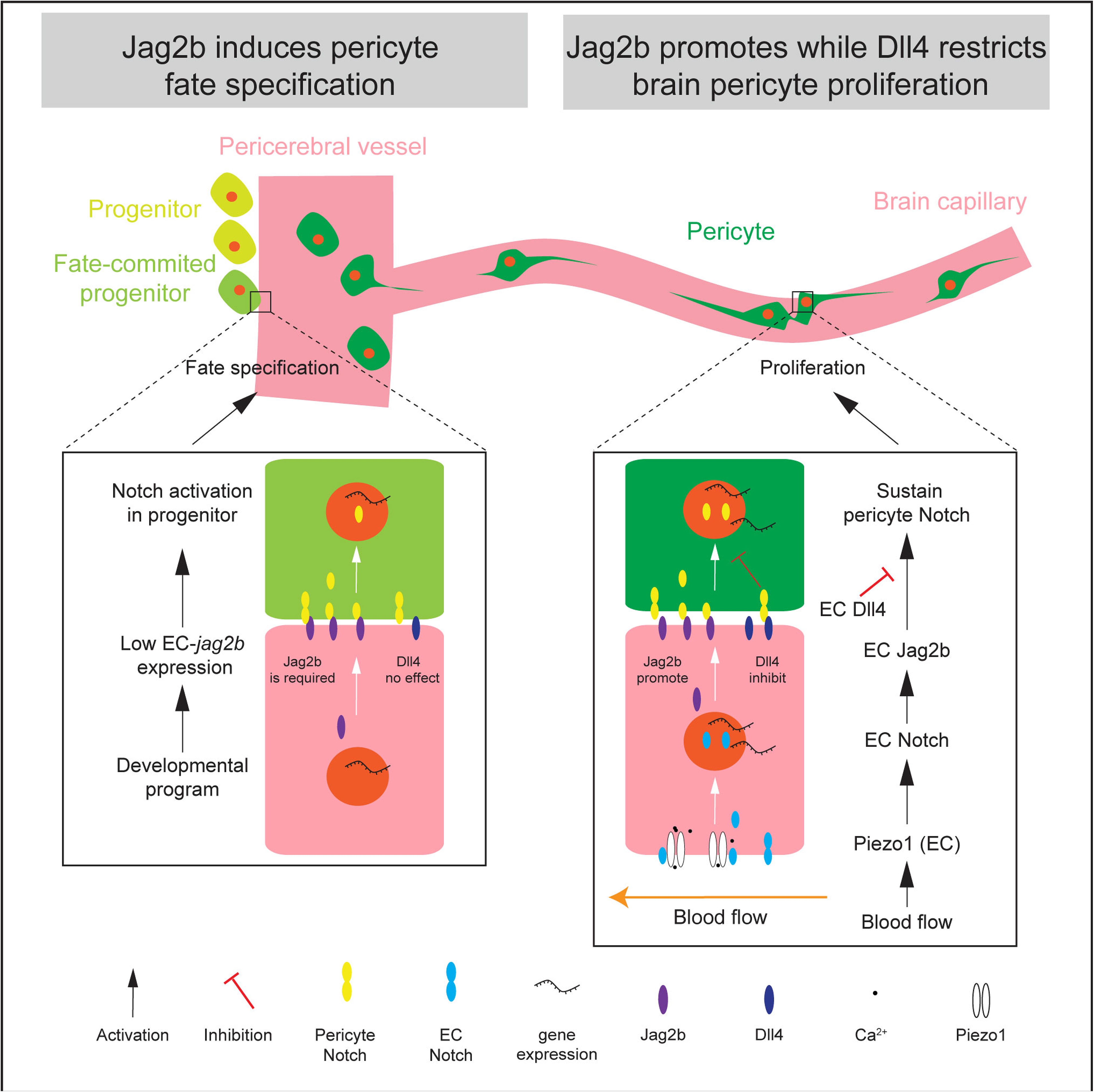

## Introduction

Pericytes are mural cells located on the abluminal surface of small blood vessels, and are key components of the neurovascular unit that support normal brain function.(Armulik *et al*, 2011; Attwell *et al*, 2016; Hill *et al*, 2015; Winkler *et al*, 2011) Loss of brain pericytes leads to multiple forms of cerebrovascular dysfunction. To maintain brain homeostasis, endothelial cells (ECs) form the blood-brain barrier (BBB) by strengthening cell-cell junctions and reducing transcytosis,(Daneman, 2012; Zhao *et al*, 2015; Zlokovic, 2008) whereas pericyte deficiency compromises BBB integrity.(Armulik *et al*, 2010; Bell *et al*, 2010; Daneman *et al*, 2010; Wang *et al*, 2014) As key regulators of blood perfusion, brain pericytes play a critical role in neurovascular coupling (NVC), a process in which neuronal activity induces localized increases in cerebral blood flow to meet metabolic demands for oxygen and glucose, while also regulating vascular tone to coordinate overall blood flow.(Alarcon-Martinez *et al*, 2020; Bell *et al*., 2010; Fernandez-Klett *et al*, 2010; Hall *et al*, 2014; Hartmann *et al*, 2021; Hartmann *et al*, 2022; Kisler *et al*, 2017; Li *et al*, 2017; Longden *et al*, 2023; Mazzoni *et al*, 2015; Peppiatt *et al*, 2006; Sun *et al*, 2026; Zambach *et al*, 2021) In addition, pericyte dysfunction has been implicated in a wide range of neurological disorders, including Alzheimer’s disease (AD), vascular dementia, and ischemic stroke.(Cai *et al*, 2017; Joutel *et al*, 1996; Liu *et al*, 2010; Nortley *et al*, 2019; Sagare *et al*, 2013; Shi *et al*, 2020; Tsao *et al*, 2021; Underly *et al*, 2017; Yang *et al*, 2022) Therefore, elucidating the mechanisms underlying brain pericyte development is essential for understanding cerebrovascular formation and for identifying potential therapeutic strategies for neurological diseases.

Brain pericyte progenitors arise from pericerebral vessels, enter the brain along nascent cerebrovascular routes, and subsequently proliferate to expand the pericyte population.^(Ando *et al*, 2016; Ando *et al*, 2019; Zi *et al*, 2024)^ Previous studies have revealed that Notch signaling is required for the fate specification of pericytes. Loss of Notch2 and Notch3 leads to defective differentiation of *pdgfrb*^+^ pericytes within the pericerebral progenitor pools during early embryogenesis.(Ando *et al*., 2019) However, the cellular source and type of Notch ligands that activate Notch receptors in pericyte progenitors during fate specification remain unknown. In addition to its role in fate specification, Notch signaling also regulates brain pericyte proliferation in a Pdgfrb-dependent manner.(Wang *et al*., 2014) Loss-of-function of Notch3 results in a marked reduction in brain pericyte number.(Wang *et al*., 2014) Hemodynamics-driven Notch signaling in ECs has been shown to activate Notch signaling in pericytes and promote their proliferation during cerebrovascular development.(Zi *et al*., 2024) However, the underlying molecular mechanisms, particularly the identity of the endothelial Notch ligand(s) mediating EC-to-pericyte signaling, remain poorly defined. Thus, the mechanisms by which Notch signaling regulates brain pericyte development require further investigation.

Zebrafish, with their transparent embryos, provide a unique advantage for *in vivo* studies of dynamic vascular cell behaviors during embryonic development.(Chávez *et al*, 2016; Lawson & Weinstein, 2002) Reporter lines labeling pericytes enable direct visualization of brain pericyte dynamics during early embryogenesis and have revealed key regulatory pathways governing this process.(Ando *et al*., 2016; Ando *et al*., 2019; Zi *et al*., 2024; Zi *et al*, 2025) Importantly, many core molecular mechanisms controlling brain pericyte development in zebrafish, including Notch signaling pathways, are conserved between zebrafish and mammals(Ando *et al*., 2016; Ando *et al*, 2021; Ando *et al*., 2019; Wang *et al*., 2014; Zi *et al*., 2024) Together, these features establish zebrafish as a powerful and efficient model for studying brain pericyte development.

In this study, we systematically investigated the roles of EC-expressed Notch ligands in brain pericyte development to elucidate the mechanisms underlying this critical regulatory pathway. Using established single-cell RNA-seq datasets, (Li *et al*, 2026; Sun *et al*., 2026) we identified Jag2b and Dll4 as the primary Notch ligands in brain ECs. Through genetic manipulation and *in vivo* imaging, we demonstrate that Jag2b, but not Dll4, is required for pericyte fate specification within pericerebral reservoirs. In addition, Jag2b promotes brain pericyte proliferation. Jag2b heterozygous mutants exhibit a significant reduction in brain pericyte numbers and decreased pericyte coverage of brain vessels. In contrast, Dll4 exerts an opposing effect on pericyte expansion. The pro-proliferative effects of hemodynamics and endothelial Notch signaling on brain pericytes are mediated by endothelial Jag2b, and the pro-expansive responses are lost in *jag2b* heterozygotes. These findings highlight a critical role for Jag2b-mediated EC-pericyte signaling in brain pericyte development.

## Results

### ECs are enriched for the Notch ligands *jag2b* and *dll4* in the brain

To investigate ligand-specific mechanisms underlying endothelial Notch regulation of brain pericyte development, we first profiled Notch ligand expression in brain ECs. Based on previously published single-cell RNA-sequencing (scRNA-seq) datasets from zebrafish brain at 6 dpf (GEO: GSE243907 and GSA: CRA030225) (Li *et al*., 2026; Sun *et al*., 2026), we analyzed 2,123 pericytes and 10,676 ECs identified using established marker genes (EC: *kdrl*, *pecam1*, *cdh5*; pericyte: *pdgfrb*, *ndufa4l2a*, *abcc9*) (Figure 1A; Figure 1—figure supplement 1A and C). These cell numbers provide representation of brain vascular cell types during early development. Additional brain cell types were annotated using canonical marker gene expression (Figure 1—figure supplement 1B and C).

**Figure 1:**
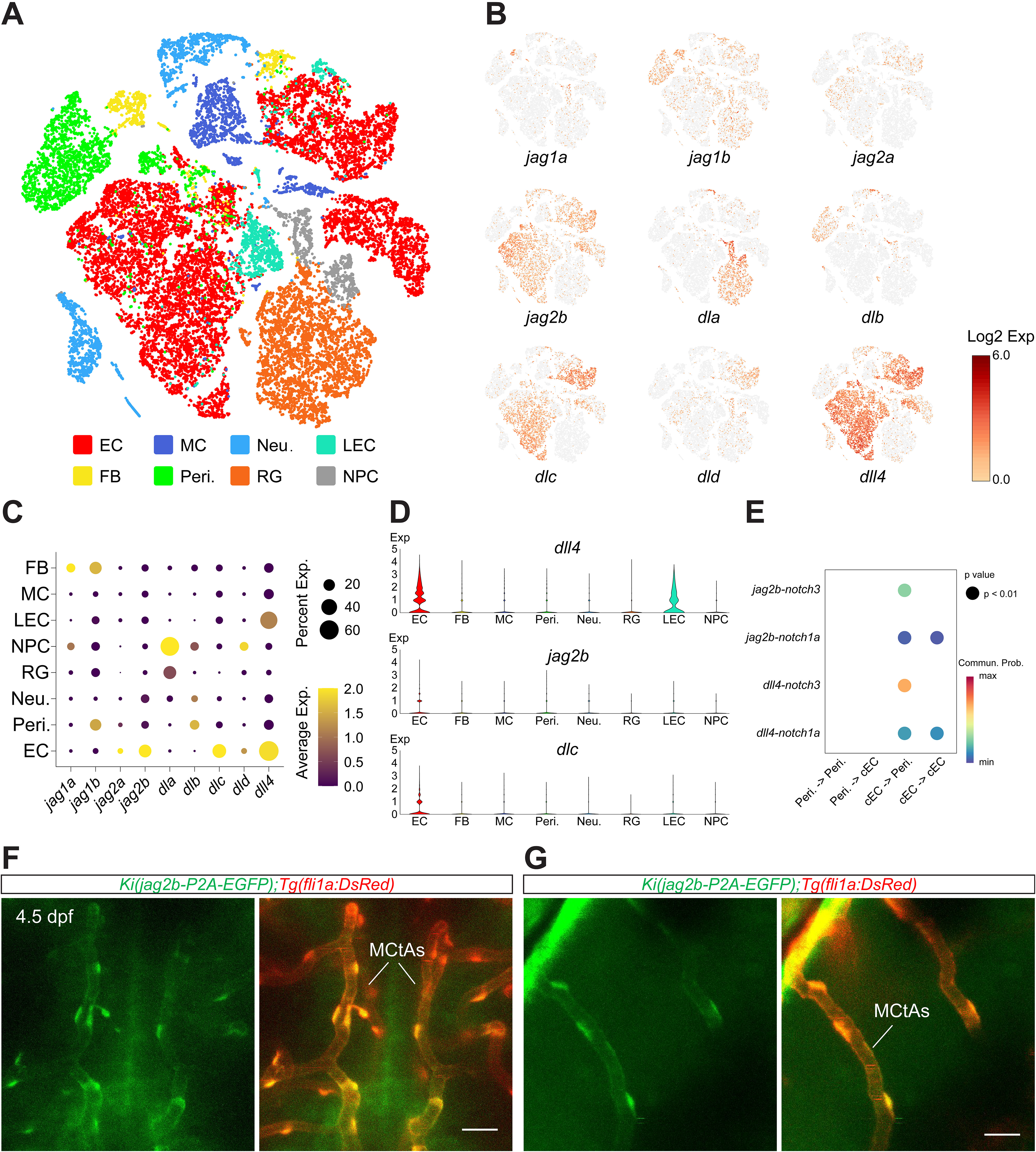
**Brain ECs are enriched for the Notch ligands *jag2b* and *dll4*** (A) t-SNE plot showing brain cells from two independent scRNA-seq datasets of 6.0- dpf zebrafish larvae. ECs and pericytes were enriched via FACS during sample preparation. Distinct cell types are color-coded: pericytes in green, ECs in red. Abbreviations: EC, endothelial cell; MC, myeloid cell; Neu., neuron; LEC, lymphatic endothelial cell; FB, fibroblast; Peri., pericyte; RG, radial glial cell; NPC, neural progenitor cell. (B) t-SNE plots showing expression levels of Notch ligand genes in 6-dpf zebrafish brains. (C) Circle plots illustrating the expression levels of Notch ligand genes across different brain cell types. (D) Violin plots showing expression levels of Notch ligand genes *dll4*, *jag2b*, and *dlc*. (E) Predicted Notch signaling interactions between brain capillary ECs and pericytes inferred by CellChat analysis. (F and G) Representative images showing *jag2b* reporter expression in brain ECs of *Ki(jag2b-EGFP);Tg(fli1a:DsRedEx)* larvae at 4.5 dpf. ECs in red, *jag2b* reporter- positive cells in green. MCtAs, mesencephalic central arteries. Images are dorsal views and partially z-projected. Scale bar: 50 μm in (F) and (G).

Notch ligands comprise two major families, Jagged and Delta-like (Kopan, 2012). We next assessed their expression across brain cell types and found that ECs are enriched for the Jagged ligand *jag2b*, and the Delta-like ligand *dll4* and *dlc* (Figure 1B– D). Notably, all three ligands are enriched in ECs, with *dll4* showing higher expression than *dlc* (Figure 1B–D). We further subclustered ECs, and identified brain capillary ECs based on the expression of *slc2a1a* and *slc7a5* (Figure 1—figure supplement 1D and E). Both *jag2b* and *dll4* are enriched in capillary ECs, whereas *dlc* is not (Figure 1— figure supplement 1F) (Li *et al*., 2026).

Focusing on capillary ECs, we performed CellChat analysis to assess EC-pericyte interactions in the brain (Figure 1—figure supplement 1G). Both Jag2b and Dll4 were predicted as EC-derived ligands potentially signaling through Notch3 and Notch1a in brain pericytes (Figure 1E; Figure 1—figure supplement 1G). Collectively, these results indicate that Jag2b and Dll4 are highly enriched in brain capillary ECs and may serve as major regulators of EC-pericyte Notch signaling.

To validate the EC-specific expression of *jag2b in vivo*, we imaged *Ki(jag2b- EGFP);Tg(fli1a:DsRedEx)* larvae, in which *EGFP* is co-expressed from the endogenous *jag2b* locus, and ECs are labeled with DsRed fluorescence (Jin *et al*, 2022). Consistent with the scRNA-seq analysis, *jag2b* reporter signal was predominantly associated with brain ECs (Figure 1F and G), confirming endothelial expression of *jag2b* in the brain. The *jag2b* reporter fluorescence was also evident in several pericerebral arteries, including the basal communicating artery (BCA), posterior communicating segment (PCS), and basilar artery (BA) (Figure 1—figure supplement 2A–D) at 4.5 dpf. In contrast, *jag2b* reporter fluorescence in the pericerebral vessels choroidal vascular plexus (CVP) and primordial hindbrain channel (PHBC) was difficult to distinguish from background (Figure 1—figure supplement 2C and D) suggesting that *jag2b* is expressed at lower levels in these vessels. Notably, *dll4* expression in brain ECs has been well documented in previous studies (Ke *et al*, 2024; Rochon *et al*, 2015).

Together, these data identify *jag2b* and *dll4* as prominent Notch ligands enriched in brain ECs. This endothelial enrichment provides a rationale for testing their roles in EC-to-pericyte Notch signaling during cerebrovascular development.

### Endothelial Jag2b is required for pericyte fate specification within pericerebral reservoirs

Brain pericytes originate from pericerebral reservoirs, where their fate is specified through an EC-dependent mechanism, and Notch signaling also plays a critical role in this process (Ando *et al*., 2016; Ando *et al*., 2019). Whether EC-derived Notch ligands regulate this process, however, remains unclear. To visualize pericyte specification *in vivo*, we performed time-lapse imaging on *Ki(pdgfrb:GAL4-VP16);Tg(4×nrUAS:GFP);Tg(fli1a:DsRedEx)* larvae (Zi *et al*., 2024), in which pericytes are labeled with GFP and ECs with DsRed. *pdgfrb* reporter-positive perivascular cells, representing newly specified pericytes, emerged around the CVP between 2.0 and 2.5 dpf (Figure 2A). Perivascular cells with no detectable *pdgfrb* reporter signal within the imaged z-stack at preceding time points progressively acquired *pdgfrb* reporter expression, became closely associated with ECs, and subsequently adopted a perivascular position along the vessel wall (Figure 2A and Figure 2—video 1), consistent with previous reports (Ando *et al*., 2016; Ando *et al*., 2019). Similar cellular behaviors were also observed during pericyte specification along other pericerebral vessels (data not shown). Within the vascular regions and developmental intervals directly examined, we did not detect a comparably prominent wave of new *pdgfrb* reporter acquisition after the initial specification period, consistent with a temporally restricted phase of early pericyte specification.

**Figure 2:**
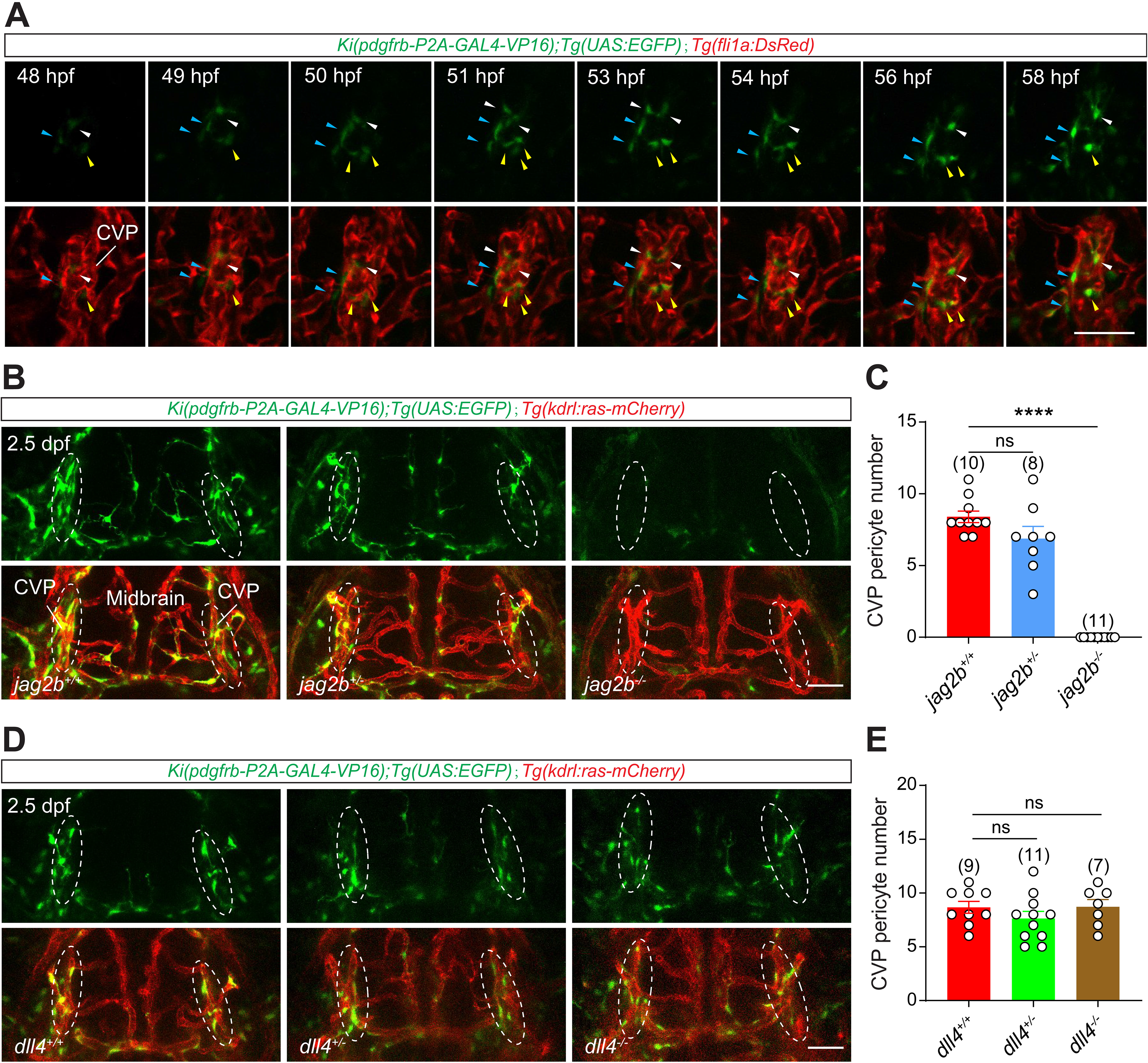
Endothelial Jag2b is required for pericyte fate specification within pericerebral reservoirs. (A) Example showing the pericyte fate specification around the pericerebral vessel CVP. (B) Representative images of the pericytes and blood vessels in the midbrain of wild- type, *jag2b* heterozygous, and *jag2b* homozygous larvae in the *Ki(pdgfrb:GAL4- VP16);Tg(4×nrUAS:GFP);Tg(kdrl:ras-mCherry)* background at 2.5 dpf. ECs in red, pericytes in green. (C) Effect of *jag2b* loss on pericyte fate specification on the CVP. (D) Representative images of the pericytes and blood vessels in the midbrain of wild- type, *dll4* heterozygous, and *dll4* homozygous larvae in the *Ki(pdgfrb:GAL4- VP16);Tg(4×nrUAS:GFP);Tg(kdrl:ras-mCherry)* background at 2.5 dpf. ECs in red, pericytes in green. (E) Effect of *dll4* loss on pericyte fate specification on the CVP. All images are dorsal views and partially z-projected. Scale bar: 50 μm for (A), (B), and (D). Images of 2.5 dpf larvae were used to quantify the number of pericytes on pericerebral vessels. Data are represented as mean ± SEM. Sample sizes (N) are indicated above the corresponding plots. Stars represent the results of unpaired two- tailed Student’s *t test* between groups (****p < 0.0001).

To determine whether EC-derived Notch ligands regulate this process, we generated *jag2b* and *dll4* mutants using CRISPR/Cas9 and quantified pericerebral pericytes at 2.5 dpf. Strikingly, *jag2b* loss abolished the appearance of newly specified pericytes in the CVP (*jag2b^-/-^* = 0 ± 0 *versus jag2b^+/+^* = 8.4 ± 0.4, decreased by 100%, p < 0.0001), whereas heterozygotes were unaffected (*jag2b^+/-^* = 6.9 ± 0.9 versus *jag2b^+/+^* = 8.4 ± 0.4, p = 0.1) (Figure 2B and C). Pericytes were also significantly reduced on the BCA/PCS/BA (*jag2b^-/-^* = 8.6 ± 0.8 *versus jag2b^+/+^* = 17.7 ± 1.0, decreased by 51%, p < 0.0001) and PHBC (*jag2b^-/-^* = 1 ± 0.2 *versus jag2b^+/+^* = 8.1 ± 1.2, decreased by 88%, p < 0.0001) (Figure 2—figure supplement 1A–C). These defects persisted at 4.5 dpf (Figure 2—figure supplement 1D and E), consistent with a persistent consequence of impaired early pericyte specification rather than a simple short developmental delay. Collectively, these data identify Jag2b as a major EC-derived signal required for efficient pericyte fate specification.

To determine whether the residual pericyte formation associated with BCA/PCS/BA observed in homozygous *jag2b* mutants remains Notch dependent, we treated mutant embryos with DAPT from 1.5 to 2.5 dpf. DAPT treatment eliminated the residual *pdgfrb*^+^ perivascular cells detected in DMSO-treated *jag2b* mutants (Figure 2—figure supplement 1F and G). These results indicate that residual pericyte formation in the absence of Jag2b remains Notch-sensitive and are consistent with a limited contribution from Jag2b-independent Notch signaling. Because DAPT inhibits Notch signaling broadly, this experiment does not identify the ligand or cellular source responsible for the residual Notch activity.

To further characterize the relationship between endothelial *jag2b* expression and pericyte specification, we examined the temporal dynamics of endogenous *jag2b* expression using *Ki(jag2b-EGFP);Tg(fli1a:DsRedEx)* larvae. *jag2b* reporter fluorescence was barely detectable in the CVP at 55 hpf (Figure 1—figure supplement 2E, G, and H), indicating relatively weak reporter expression in this pericerebral progenitor niche during the period of pericyte specification. Similarly, *jag2b* reporter fluorescence was also difficult to distinguish from background in other pericerebral vessels at 55 hpf (Figure 1—figure supplement 2E and F), indicating weak reporter activity in these vascular regions at this stage. Moreover, pericyte specification was not significantly altered in either *jag2b* heterozygous mutants or EC-*jag2b*-overexpressing embryos (Figure 2B and C; Figure 2—figure supplement 1A–C, H, and I). Thus, although Jag2b is required for efficient pericyte specification, increasing its endothelial expression is not sufficient to enhance this process. Together with the weak reporter expression observed during the specification period, these findings are consistent with a permissive rather than strongly instructive role for endothelial Jag2b.

In contrast, *dll4* mutants showed no defect in pericyte specification (*dll4^-/-^* = 8.7 ± 0.7 *versus dll4^+/+^* = 8.7 ± 0.6, p = 0.96) (Figure 2D and E), indicating that Dll4 is not detectably required for this process under these conditions.

Together, these results identify endothelial Jag2b as a major component of the signaling environment required for efficient pericyte fate specification within pericerebral reservoirs during a restricted developmental window.

### Endothelial Jag2b promotes brain pericyte proliferation and increases pericyte coverage of brain vessels

After entering the brain, pericytes expand their population primarily through *in situ* proliferation (Figure 3—video 1), a process regulated by endothelial-to-pericyte Notch signaling (Zi *et al*., 2024). We therefore asked which endothelial Notch ligand(s) promote this post-specification expansion. Because homozygous *jag2b* mutants exhibit a severe early defect in pericyte specification, whereas *jag2b* heterozygous mutants show no significant defect in this process at 2.5 dpf (Figure 2B and C), we used heterozygous mutants to examine post-specification functions of Jag2b while minimizing the confounding effect of the early specification phenotype.

**Figure 3:**
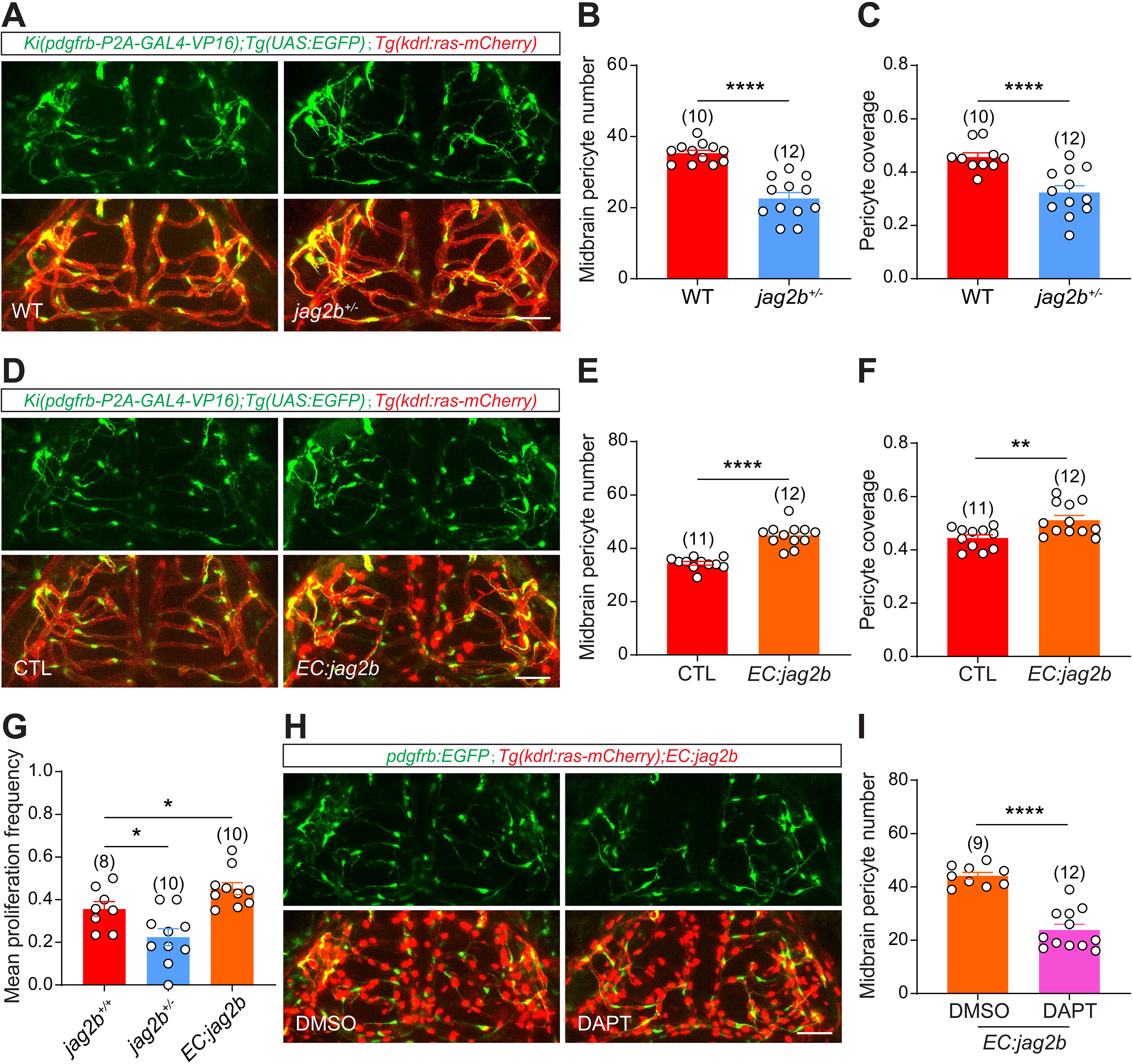
Endothelial Jag2b promotes brain pericyte proliferation and increases pericyte coverage of brain vessels. (A) Representative images of the pericytes and blood vessels in the midbrain of wild- type and *jag2b* heterozygous larvae in the *Ki(pdgfrb:GAL4- VP16);Tg(4×nrUAS:GFP);Tg(kdrl:ras-mCherry)* background at 4.5 dpf. ECs in red, pericytes in green. (B) Effect of *jag2b* loss on midbrain pericyte number. (C) Effect of *jag2b* loss on pericyte coverage of blood vessels in the midbrain. (D) Representative images of the pericytes and blood vessels in the midbrain of control and EC-specific *jag2b* overexpression larvae in the *Ki(pdgfrb:GAL4- VP16);Tg(4×nrUAS:GFP);Tg(kdrl:ras-mCherry)* background at 4.5 dpf. ECs and nucleus of *jag2b*-overexpressing cells in red, pericytes in green. (E) Effect of EC-specific *jag2b* overexpression on pericyte number in the midbrain. (F) Effect of EC-specific *jag2b* overexpression on pericyte coverage of blood vessels in the midbrain. (G) Effects of *jag2b* loss and EC-specific *jag2b* overexpression on pericyte proliferation frequency in the midbrain. (H) Representative images of the pericytes and blood vessels in the midbrain of *Ki(pdgfrb:GAL4-VP16);Tg(4×nrUAS:GFP);Tg(kdrl:ras-mCherry);Tg(kdrl:H2B- mScarlet-P2A-jag2b)* larvae with DMSO or DAPT treatment at 4.5 dpf. ECs and nucleus of *jag2b*-overexpressing cells in red, pericytes in green. (I) Effect of DAPT treatment on midbrain pericyte number in larvae with EC-specific *jag2b* overexpression. All images are dorsal views and partially z-projected. Scale bar: 50 μm in (A), (D), and (H). Images of 4.5 dpf larvae were used to quantify midbrain pericyte number and pericyte coverage of midbrain vessels, time-lapse images from 3.0-3.5 dpf were used to quantify the mean proliferation frequency of midbrain pericytes. Data are represented as mean ± SEM. Sample sizes (N) are indicated above the corresponding plots. Stars represent the results of unpaired two-tailed Student’s t test between groups (*p < 0.05, **p < 0.01, ****p < 0.0001).

At 4.5 dpf, *jag2b*^+/-^ larvae showed a significant reduction in midbrain pericyte number (*jag2b*^+/-^ = 22.6 ± 1.7 versus WT = 35.1 ± 1.0, decreased by 36%, p < 0.0001) (Figure 3A and B), compared with the wild-type (WT). Given reduced vascular density in homozygous *jag2b* mutants (Figure 3—figure supplement 1A and C), we additionally quantified pericyte coverage to brain vessels as previously described (Xu *et al*, 2017), to confirm the reduction in pericyte density (*jag2b*^+/-^ = 0.32 ± 0.03 versus WT = 0.46 ± 0.02, decreased by 30%, p < 0.0001) (Figure 3C). The reduced phenotype persisted into adulthood (Figure 3—figure supplement 1E), consistent with a lasting consequence of the developmental defect.

Brain pericyte population expansion during early development reflects both ingress of pericerebral pericytes and *in situ* proliferation (Zi *et al*., 2024). Long-term time-lapse imaging (3.0-3.5 dpf) revealed that *jag2b*^⁺/⁻^ larvae exhibited reduced pericyte proliferation (*jag2b^+/-^* = 0.22 ± 0.04 *versus* WT = 0.36 ± 0.03, decreased by 39%, p < 0.05) (Figure 3G), while pericyte ingress was unchanged (Figure 3—figure supplement 1D). Together, these results indicate that Jag2b primarily regulates the *in situ* proliferation of brain pericytes during brain vascular development rather than the ingress of pericerebral pericytes into the brain.

To test sufficiency, we generated a *Tg(kdrl:H2B-mScarlet-P2A-jag2b)* transgenic line for EC-specific *jag2b* overexpression (Figure 3—figure supplement 1B).

Importantly, endothelial expression of the P2A-linked *jag2b* transgene partially rescued the brain pericyte phenotype of homozygous *jag2b* mutants (*EC:jag2b+jag2b^-/-^* = 11.4 ± 0.9 *versus jag2b^-/-^* = 7.1 ± 0.6, increased by 61%, p < 0.01) (Figure 3—figure supplement 1F and G), confirming that the P2A-linked *jag2b* transgene retains substantial biological activity.

In otherwise WT larvae, EC-specific *jag2b* overexpression significantly increased midbrain pericyte number (*EC:jag2b* = 44.8 ± 1.2 *versus* WT control = 34.4 ± 0.7, increased by 30%, p < 0.0001) and pericyte coverage of midbrain vessels (*EC:jag2b* = 0.51 ± 0.02 *versus* WT control = 0.44 ± 0.01, increased by 16%, p < 0.01) (Figure 3D– G; Figure 3—figure supplement 1D). Consistently, time-lapse imaging showed an increased brain pericyte proliferation frequency in EC-*jag2b*-overexpressing larvae (*EC:jag2b* = 0.45 ± 0.03 *versus* WT control = 0.36 ± 0.03, increased by 25%, p < 0.05) (Figure 3G), whereas pericyte ingress was not significantly affected (Figure 3—figure supplement 1D). Consistent with these findings, analysis of proliferation-marker expression further supported the pro-expansive effect of endothelial Jag2b (Figure 3— figure supplement 1H). Together, these loss- and gain-of-function analyses indicate that endothelial Jag2b promotes brain pericyte population expansion primarily by enhancing *in situ* proliferation.

We next asked whether the pro-expansive effect of endothelial Jag2b requires Notch signaling. DAPT was applied from 3.5 to 4.5 dpf, after the major period of initial pericyte specification, to preferentially assess the role of Notch signaling during subsequent pericyte population expansion. Under these conditions, Notch inhibition markedly reduced the increased brain pericyte number induced by EC-specific *jag2b* overexpression (DAPT = 23.8 ± 2.1 versus DMSO control = 44.1 ± 1.3, decreased by 46%, p < 0.0001) (Figure 3H and I), supporting a requirement for Notch activity in Jag2b-driven pericyte expansion.

Because PDGFRβ signaling is also required for brain pericyte proliferation, we further examined whether PDGFRβ activity contributes to the proliferative response associated with endothelial Jag2b overexpression. Treatment with the PDGFRβ inhibitor AG1295 during the 3.0–3.5 dpf time-lapse imaging period significantly reduced brain pericyte proliferation in EC-*NICD*-overexpressing larvae (AG1295 = 0.21 ± 0.01 versus DMSO control = 0.57 ± 0.06, decreased by 63%, p < 0.001) (Figure 3—figure supplement 1I), consistent with our previous finding that Notch-dependent pericyte proliferation requires PDGFRβ signaling.^11^ We next asked whether the increase in brain pericytes induced by endothelial *jag2b* overexpression similarly depends on PDGFRβ activity. AG1295 treatment also significantly attenuated the EC- *jag2b*-overexpression-induced increase in brain pericyte proliferation frequency (AG1295 = 0.33 ± 0.03 versus DMSO control = 0.51 ± 0.05, decreased by 35%, p < 0.01) (Figure 3—figure supplement 1J). Together, these results indicate that intact PDGFRβ signaling is required to sustain the enhanced pericyte proliferative response induced by endothelial *jag2b* overexpression.

Together, these findings identify endothelial Jag2b as a positive regulator of post-specification brain pericyte expansion, acting primarily by promoting Notch-dependent *in situ* proliferation that requires intact PDGFRβ activity.

### Endothelial Dll4 negatively regulates brain pericyte proliferation

We next tested whether Dll4 regulates brain pericyte expansion. At 4.5 dpf, *dll4*^⁻/⁻^ larvae exhibited increased midbrain pericyte number (*dll4^-/-^* = 58.8 ± 3.0 *versus* WT = 37.1 ± 0.9, increased by 58%, p < 0.0001), vessel segment number (*dll4^-/-^* = 105.7 ± 4.7 *versus* WT = 82.0 ± 2.0, increased by 29%, p < 0.001), endothelial tip cell (ETC) number (*dll4^-/-^* = 15.3 ± 0.7 *versus* WT = 4.9 ± 0.4, increased by 212%, p < 0.0001). Because loss of dll4 also increased the extent of the cerebrovascular network, we additionally quantified pericyte coverage relative to the vascular network. Pericyte coverage remained significantly increased in *dll4^-/-^* larvae (*dll4^-/-^* = 0.56 ± 0.02 *versus* WT = 0.45 ± 0.01, increased by 24%, p < 0.001) (Figure 4A, B, and E; Figure 4—figure supplement 1A, C, and D). Thus, the increase in pericyte abundance cannot be explained solely by the accompanying increase in brain vessel segment number. To directly assess whether Dll4 affects pericyte population expansion through changes in pericyte proliferation, we performed time-lapse imaging from 3.0 to 3.5 dpf. Loss of *dll4* significantly increased brain pericyte proliferation frequency (*dll4^-/-^* = 0.47 ± 0.02 *versus* WT control = 0.34 ± 0.03, increased by 38%, p < 0.01) (Figure 4G), whereas pericyte ingress into the brain was not detectably altered (Figure 4—figure supplement 1E). These findings indicate that Dll4 negatively regulates brain pericyte population expansion primarily by restraining pericyte proliferation rather than by altering pericyte ingress.

**Figure 4:**
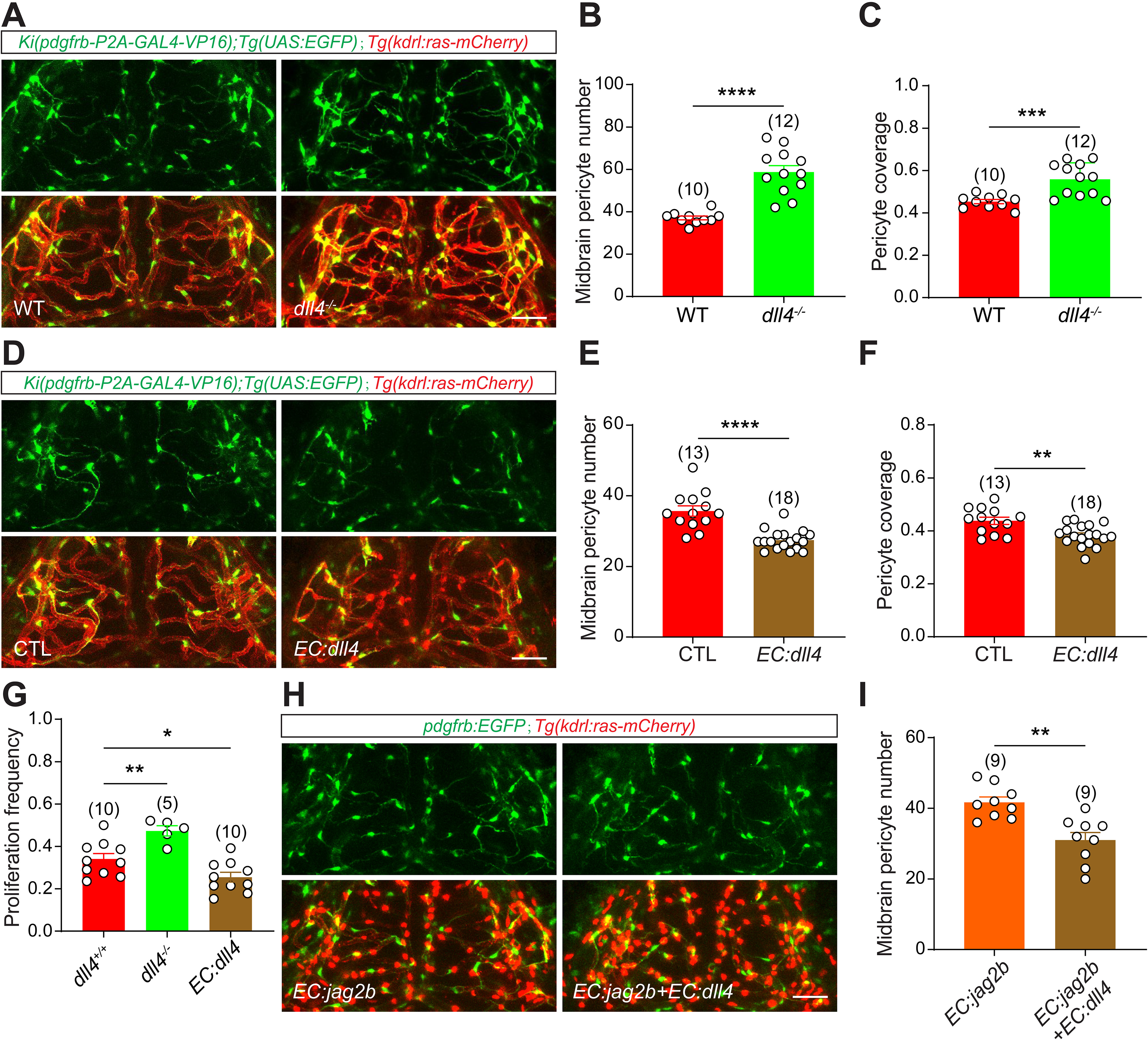
Endothelial Dll4 negatively regulates brain pericyte proliferation. (A) Representative images of the pericytes and blood vessels in the midbrain of wild- type and *dll4* homozygous larvae in the *Ki(pdgfrb:GAL4- VP16);Tg(4×nrUAS:GFP);Tg(kdrl:ras-mCherry)* background at 4.5 dpf. ECs in red, pericytes in green. (B) Effect of *dll4* loss on midbrain pericyte number. (C) Representative images of the pericytes and blood vessels in the midbrain of control and EC-specific *dll4* overexpression larvae in the *Ki(pdgfrb:GAL4- VP16);Tg(4×nrUAS:GFP);Tg(kdrl:ras-mCherry)* background at 4.5 dpf. ECs and nucleus of *dll4*-overexpressing cells in red, pericytes in green. (D) Effect of EC-specific *dll4* overexpression on midbrain pericyte number. (E) Effect of *dll4* loss on pericyte coverage of blood vessels in the midbrain. (F) Effect of EC-specific *dll4* overexpression on pericyte coverage of blood vessels in the midbrain. (G) Effects of *dll4* loss and EC-specific *dll4* overexpression on midbrain pericyte proliferation frequency. (H) Representative images of midbrain pericytes and blood vessels in EC-*jag2b*- overexpressing larvae and larvae with simultaneous endothelial overexpression of *jag2b* and *dll4* in the *Ki(pdgfrb:GAL4-VP16);Tg(4×nrUAS:GFP);Tg(kdrl:ras- mCherry)* background at 4.5 dpf. ECs and the nuclei of *jag2b*/*dll4*-overexpressing cells in red, pericytes in green. (I) Effect of EC-*dll4* overexpression on midbrain pericyte number in EC-*jag2b*- overexpressing larvae. All images are dorsal views and partially z-projected. Scale bar: 50 μm in (A), (C), and (H). Images of 4.5 dpf larvae were used to quantify midbrain pericyte number and pericyte coverage of midbrain vessels, time-lapse images from 3.0-3.5 dpf to quantify the mean proliferation frequency of midbrain pericytes. Data are represented as mean ± SEM. Sample sizes (N) are indicated above the corresponding plots. Stars represent the results of unpaired two-tailed Student’s t test between groups (*p < 0.05, **p < 0.01, ***p < 0.001, ****p < 0.0001).

To determine whether increased endothelial Dll4 activity produces the reciprocal phenotype, we generated a *Tg(kdrl:H2B-mScarlet-P2A-dll4)* transgenic line, in which EC nuclei are labeled with H2B-mScarlet and *dll4* is overexpressed specifically in ECs (Figure 4—figure supplement 1B). *EC:dll4* larvae showed reduced midbrain pericyte number (*EC:dll4* = 27.4 ± 0.7 *versus* WT control = 35.7 ± 1.5, decreased by 23%, p < 0.0001), vessel segment number (*EC:dll4* = 71.9 ± 1.6 *versus* WT control = 81.5 ± 2.0, decreased by 12%, p < 0.001), and pericyte coverage of midbrain vessels (*EC:dll4* = 0.38 ± 0.01 *versus* WT control = 0.44 ± 0.01, decreased by 14%, p < 0.01) (Figure 4C, D, and F; Figure 4—figure supplement 1A and C). In addition, time-lapse imaging (3.0- 3.5 dpf) confirmed suppressed pericyte proliferation (*EC:dll4* = 0.25 ± 0.02 *versus* WT control = 0.34 ± 0.03, p = 0.02) (Figure 4G), without altering ingress (Figure 4—figure supplement 1E), in *EC:dll4* larvae. Thus, endothelial Dll4 inhibits the expansion of the brain pericyte population by limiting pericyte proliferation during development.

Because Jag2b promotes pericyte proliferation whereas Dll4 suppresses it, we next asked whether Dll4 functionally opposes Jag2b-dependent pericyte expansion. Endothelial *jag2b* overexpression increased brain pericyte number, whereas simultaneous endothelial overexpression of *dll4* significantly suppressed this Jag2b- induced increase (*EC:jag2b*+*EC:dll4* = 31.0 ± 2.2 versus *EC:jag2b* = 41.7 ± 1.5; decreased by 26%, p < 0.0001) (Figure 4H and I). Thus, elevated endothelial Dll4 activity can counteract the pro-expansive effect of Jag2b.

We further examined the genetic interaction between *jag2b* and *dll4*. Partial CRISPR-mediated disruption of *jag2b* significantly reduced the elevated brain pericyte number observed in *dll4* mutant larvae (Cas9+*jag2b* sgRNA = 3.4 ± 1.2 versus Cas9 Control = 50.7 ± 2.5, decreased by 93%, p < 0.01) (Figure 4—figure supplement 1F and G). Although this experiment cannot fully distinguish effects on initial pericyte specification from those on subsequent population expansion, it provides additional genetic evidence that the increased pericyte population associated with *dll4* loss depends, at least in part, on Jag2b function.

Together, these findings demonstrate that endothelial Jag2b and Dll4 exert functionally opposing effects on brain pericyte population expansion, with Jag2b promoting and Dll4 restraining pericyte proliferation during cerebrovascular development.

### Endothelial Jag2b mediates the pro-expansive effects of EC-intrinsic Notch signaling on brain pericytes

Given that EC-intrinsic Notch signaling promotes brain pericyte proliferation by triggering EC-to-pericyte signaling (Zi *et al*., 2024), we next asked whether Jag2b mediates this interaction.

To determine whether EC-intrinsic Notch activity regulates *jag2b* expression, we generated a *Tg(kdrl:tetoff-H2B-mScarlet-P2A-NICD)* line (Figure 5—figure supplement 1A), in which Notch signaling is activated specifically in ECs. Functional activation of endothelial Notch signaling was confirmed using the *in vivo* Notch reporter line *Tg(Tp1:d2GFP)*, which showed increased reporter activity in *NICD*- expressing ECs (Figure 5—figure supplement 1C and D) (Zi *et al*., 2024). We then used the *Ki(jag2b-EGFP)* line to monitor endogenous *jag2b* expression. Imaging of *Tg(kdrl:tetoff-H2B-mScarlet-P2A-NICD);Ki(jag2b-EGFP);Tg(fli1a:DsRedEx)* larvae revealed increased *jag2b* reporter expression in *NICD*-expressing midbrain ECs (Figure 5A and B). Consistent with this observation, qPCR analysis further showed increased *jag2b* expression in brain ECs following endothelial *NICD* overexpression (Figure 5C). Together, these findings indicate that activation of endothelial Notch signaling positively regulates *jag2b* expression.

**Figure 5:**
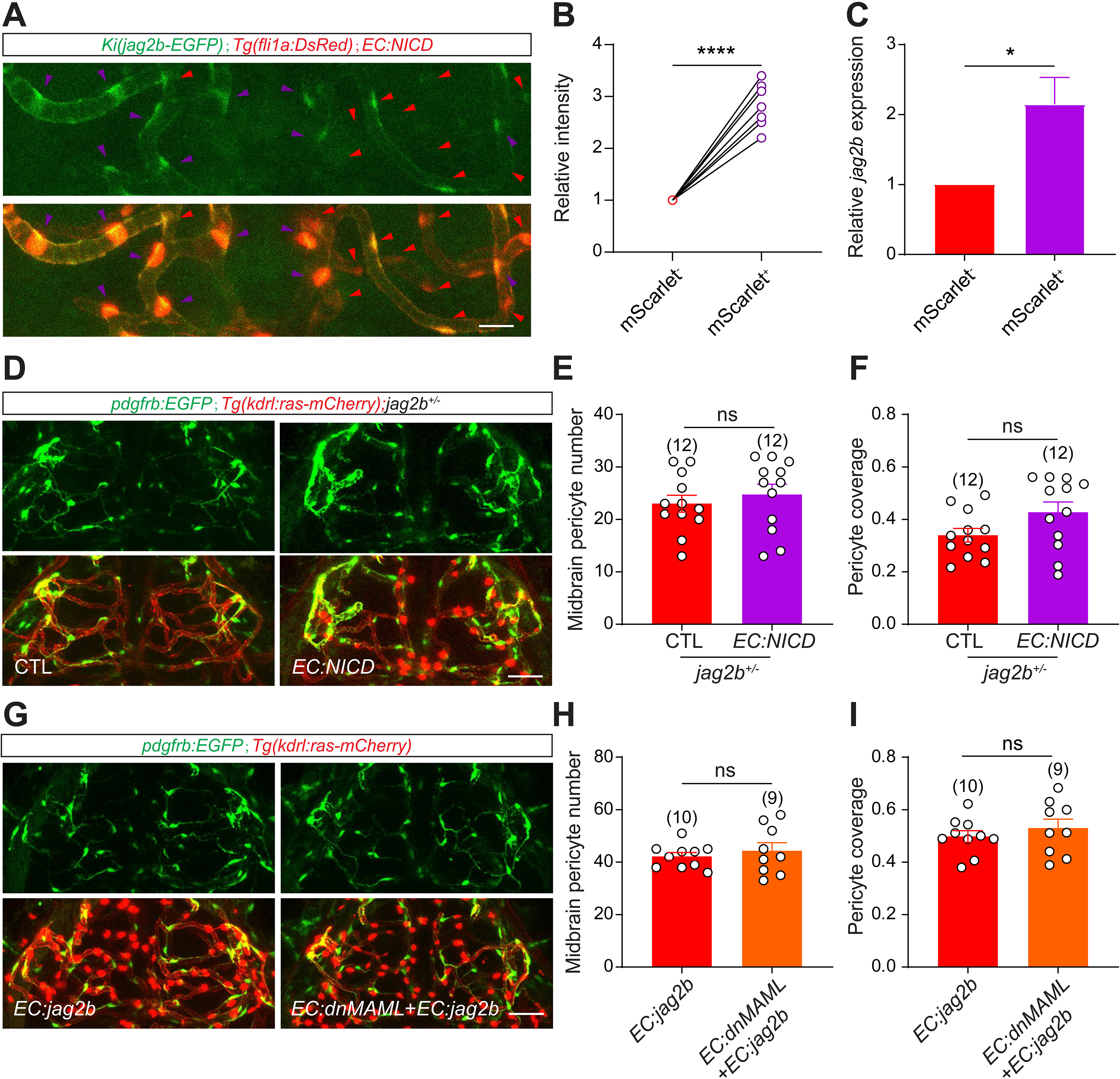
Endothelial Jag2b mediates the pro-expansive effects of EC-intrinsic Notch signaling on brain pericytes. (A) Representative images of the blood vessels and endogenous *jag2b* expression in the midbrain of *Ki(jag2b-EGFP);Tg(fli1a:DsRedEx);Tg(kdrl:tetoff-H2B-mScarlet-P2A-NICD)* larvae at 4.5 dpf. ECs and the nucleus of *NICD*-overexpressing cells in red, the cells expressing *jag2b* reporter signals in green. (B) Effect of endothelial Notch activation on endogenous *jag2b* expression in midbrain ECs. (C) Relative expression of *jag2b* in FACS-isolated EC-*NICD* brain ECs compared with control brain ECs at 6.0 dpf. (D) Representative images of the pericytes and blood vessels in the midbrain of control and EC-specific *NICD* overexpression larvae in the *Ki(pdgfrb:GAL4- VP16);Tg(4×nrUAS:GFP);Tg(kdrl:ras-mCherry); jag2b^+/-^* background at 4.5 dpf. ECs and nuclei of *NICD*-overexpressing cells in red, pericytes in green. (E) Effect of EC-specific *NICD* overexpression on midbrain pericyte number in *jag2b* heterozygous larvae. (F) Effect of EC-specific *NICD* overexpression on pericyte coverage of midbrain vessels in *jag2b* heterozygous larvae. (G) Representative images of midbrain pericytes and blood vessels in EC-*jag2b* larvae and *EC-dnMAML;EC-jag2b* larvae at 4.5 dpf. ECs and nucleus of *jag2b*/*dnMAML*- overexpressing cells in red, pericytes in green. (H) Effect of EC-*dnMAML* overexpression on midbrain pericyte number in EC-*jag2b*- overexpressing larvae. (I) Effect of EC-*dnMAML* overexpression on pericyte coverage of midbrain vessels in EC-*jag2b*-overexpressing larvae. All images are dorsal views and partially z-projected. Scale bar: 50 μm in (A), (D), and (G). Images of 4.5 dpf larvae were used to quantify midbrain pericyte number and pericyte coverage of midbrain vessels. Data are represented as mean ± SEM. N values are shown above the corresponding plots. Stars represent the results of unpaired two- tailed Student’s t test between groups (*p < 0.05, ***p < 0.001, ****p < 0.0001).

We next tested whether Jag2b is required for endothelial Notch-dependent pericyte expansion. Consistent with previous reports (Zi *et al*., 2024), EC-specific *NICD* overexpression increased midbrain pericyte number (*EC:NICD* = 53.0 ± 3.1 *versus* control = 35.2 ± 1.5, increased by 51%, p < 0.0001) and pericyte coverage of midbrain vessels (*EC:NICD* = 0.85 ± 0.04 *versus* control = 0.46 ± 0.02, increased by 85%, p < 0.0001) (Figure 5—figure supplement 1F–H), accompanied by reduced cerebrovascular growth (Figure 5—figure supplement 1E). However, in *jag2b^⁺/⁻^* mutants, endothelial Notch activation failed to increase midbrain pericyte number (*EC:NICD* = 24.8 ± 2.0 *versus* control = 23.0 ± 1.6, p = 0.50) or vascular pericyte coverage (*EC:NICD* = 0.43 ± 0.04 *versus* control = 0.34 ± 0.03, p = 0.08) (Figure 5D–F), indicating that Jag2b is required for EC-Notch-induced brain pericyte expansion.

To test sufficiency, we examined whether EC-specific *jag2b* overexpression could rescue the defects caused by endothelial Notch inhibition. We generated a *Tg(kdrl:tetoff-H2B-mScarlet-P2A-dnMAML)* line to attenuate canonical MAML- dependent Notch transcription in ECs through a dominant-negative mechanism (Figure 5—figure supplement 1B) (Zi *et al*., 2024), which reduced midbrain pericyte number (*EC:dnMAML* = 28.6 ± 1.0 *versus* control = 37.9 ± 1.6, decreased by 25%, p < 0.0001) and pericyte coverage of midbrain vessels (*EC:dnMAML* = 0.34 ± 0.01 *versus* control = 0.44 ± 0.01, decreased by 23%, p < 0.0001) (Figure 5—figure supplement 1I–K), while cerebrovascular growth remained relatively preserved (Figure 5—figure supplement 1E). To test whether elevated endothelial Jag2b can bypass attenuation of EC-intrinsic Notch signaling, we compared EC*-jag2b*-overexpressing larvae in the presence or absence of EC-*dnMAML*. EC-*dnMAML* did not significantly reduce midbrain pericyte number (*EC:jag2b* + *EC:dnMAML* = 42.6 ± 2.3 versus *EC:jag2b* = 43.0 ± 1.8, p = 0.90) or the pericyte coverage of midbrain vessels (*EC:jag2b* + *EC:dnMAML* = 0.49 ± 0.02 versus *EC:jag2b* = 0.50 ± 0.01, p = 0.65) in the EC-*jag2b*- overexpressing background (Figure 5G–I). These results are consistent with elevated endothelial Jag2b bypassing the requirement for full EC-intrinsic Notch activity in maintaining brain pericyte expansion.

Taken together, these findings demonstrate that endothelial Jag2b acts downstream of EC-intrinsic Notch signaling to drive the expansion of the brain pericyte population.

### Endothelial Jag2b mediates the pro-expansive effects of hemodynamics on brain pericytes

We previously reported that hemodynamics promote brain pericyte proliferation through activation of EC-intrinsic Notch signaling (Zi *et al*., 2024). Here, we asked whether Jag2b participates in the hemodynamics–Piezo1–EC-Notch pathway that promotes brain pericyte expansion.

Because the mechanosensitive ion channel Piezo1 transduces blood flow signals in ECs to promote brain pericyte proliferation (Zi *et al*., 2024), we tested whether Jag2b functions within this pathway. Consistent with prior findings (Zi *et al*., 2024), pharmacological activation of Piezo1 with Yoda1 increased pericyte coverage of midbrain vessels (Yoda1 = 0.62 ± 0.04 *versus* DMSO control = 0.43 ± 0.02, increased by 44%, p < 0.0001), without significantly affecting brain pericyte number (Figure 6— figure supplement 1A–C). However, this effect was abolished in *jag2b^⁺/⁻^* mutants (Yoda1 = 0.32 ± 0.02 *versus* DMSO control = 0.36 ± 0.02, p = 0.25) (Figure 6A–C), indicating that Jag2b is required for Piezo1-driven brain pericyte expansion.

**Figure 6:**
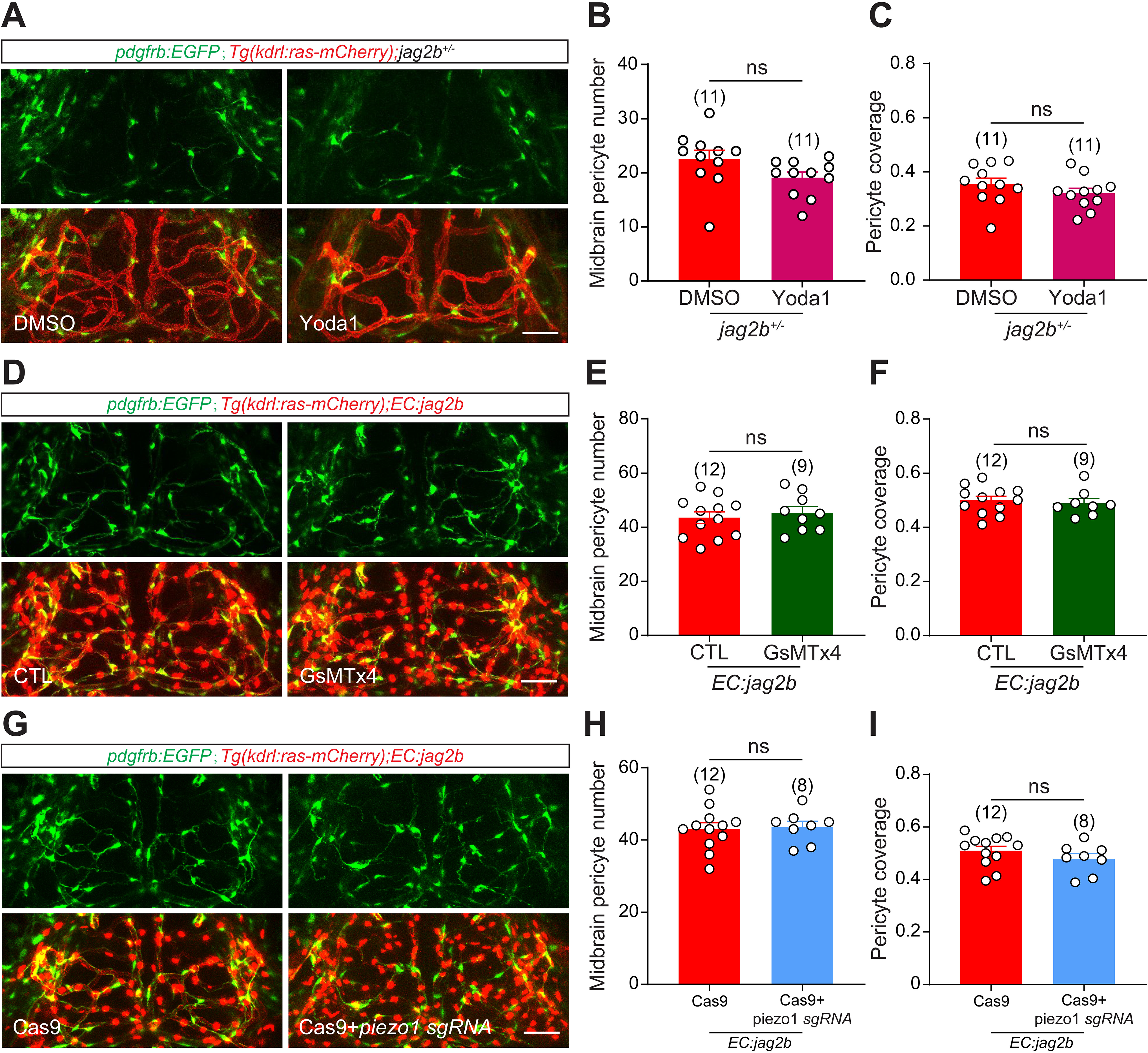
Endothelial Jag2b mediates the pro-expansive effects of hemodynamics on brain pericytes. (A) Representative images of the pericytes and blood vessels in the midbrain of DMSO and Yoda1 treated larvae in the *Ki(pdgfrb:GAL4- VP16);Tg(4×nrUAS:GFP);Tg(kdrl:ras-mCherry); jag2b^+/-^* background at 4.5 dpf. ECs in red, pericytes in green. (B) Effect of Yoda1 application on midbrain pericyte number in *jag2b* heterozygous larvae. (C) Effect of Yoda1 application on pericyte coverage of midbrain vessels in *jag2b* heterozygous larvae. (D) Representative images of the pericytes and blood vessels in the midbrain of control and GsMTx4 treated larvae in the *Ki(pdgfrb:GAL4- VP16);Tg(4×nrUAS:GFP);Tg(kdrl:ras-mCherry); Tg(kdrl:H2B-mScarlet-P2A-jag2b)* background at 4.5 dpf. ECs and nucleus of *jag2b*-overexpressing cells in red, pericytes in green. (E) Effect of GsMTx4 application on midbrain pericyte number in the larvae with EC- specific *jag2b* overexpression. (F) Effect of GsMTx4 application on pericyte coverage of midbrain vessels in the larvae with EC-specific *jag2b* overexpression. (G) Representative images of midbrain pericytes and blood vessels in EC-*jag2b*- overexpressing larvae injected with Cas9 alone or Cas9 together with *piezo1* sgRNA in the *Ki(pdgfrb:GAL4-VP16);Tg(4×nrUAS:GFP);Tg(kdrl:ras-mCherry)* background at 4.5 dpf. ECs in red, pericytes in green. (H) Effect of CRISPR-mediated disruption of *piezo1* on midbrain pericyte number in EC-*jag2b*-overexpressing larvae. (I) Effect of CRISPR-mediated disruption of *piezo1* on pericyte coverage of midbrain vessels in EC-*jag2b*-overexpressing larvae. All images are dorsal views and partially z-projected. Scale bar: 50 μm in (A), (D), and (G). Images of 4.5 dpf larvae were used to quantify midbrain pericyte number and pericyte coverage of midbrain vessels. Data are represented as mean ± SEM. Sample sizes (N) are indicated above the corresponding plots. Stars represent the results of unpaired two-tailed Student’s t test between groups.

We next inhibited Piezo1 using GsMTx4 (Bae *et al*, 2011). As expected, Piezo1 inhibition reduced midbrain pericyte number (GsMTx4 = 32.8 ± 1.1 *versus* Control = 37.2 ± 1.8, decreased by 15%, p < 0.05) and pericyte coverage of midbrain vessels (GsMTx4 = 0.37 ± 0.01 *versus* Control = 0.48 ± 0.02, decreased by 23%, p < 0.0001) (Figure 6—figure supplement 1D–F), consistent with piezo1 mutant phenotypes (Zi *et al*., 2024). Notably, in EC-specific *jag2b*-overexpression larvae, GsMTx4 treatment no longer reduced midbrain pericyte number (GsMTx4 = 45.3 ± 2.3 *versus* Control = 43.5 ± 2.1, p = 0.57) or pericyte coverage of midbrain vessels (GsMTx4 = 0.49 ± 0.02 *versus* Control = 0.50 ± 0.01, p = 0.65) (Figure 6D–F), consistent with Jag2b functioning downstream of Piezo1.

To further validate the role of Piezo1 using a gene-specific approach, we generated *piezo1* crispants by co-injection of Cas9 and *piezo1* sgRNA. CRISPR-mediated disruption of *piezo1* reduced both brain pericyte number (*piezo1* crispant = 22.2 ± 1.7 *versus* Control = 34.7 ± 1.4, decreased by 36%, p < 0.0001) and pericyte coverage of brain vessels (*piezo1* crispant = 0.26 ± 0.02 *versus* Control = 0.46 ± 0.02, decreased by 43%, p < 0.0001) (Figure 6—figure supplement 1G–I), recapitulating the phenotype of the stable piezo1 knockout line (Zi *et al*., 2024). By contrast, EC-specific *jag2b* overexpression rescued these pericyte defects in *piezo1* crispants, with no significant reduction in either brain pericyte number (*piezo1* crispant = 43.6 ± 1.6 *versus* Control = 43.1 ± 1.7, p = 0.83) or pericyte coverage (*piezo1* crispant = 0.48 ± 0.02 *versus* Control = 0.51 ± 0.02, p = 0.26) (Figure 6G–I).

Together with our previous endothelial-specific Piezo1 loss-of-function data (Zi *et al*., 2024), these results provide genetic evidence consistent with Jag2b functioning downstream of the endothelial Piezo1–Notch axis in regulating brain pericyte expansion.

## Discussion

Here, we define stage-dependent, ligand-specific roles of endothelial Notch ligands in brain pericyte development. During the early specification period, Jag2b is required for efficient emergence of *pdgfrb^+^* pericyte progenitors from pericerebral vascular niches, whereas increasing endothelial Jag2b is not sufficient to enhance specification, consistent with a permissive role. After specification, Jag2b promotes Notch-dependent pericyte proliferation and population expansion, whereas Dll4 exerts an opposing effect by restraining proliferation. These findings reveal that endothelial Notch ligands regulate distinct phases of brain pericyte development through functionally different, rather than simply redundant, ligand inputs.

The hemodynamics–EC Notch–pericyte regulatory axis is known to regulate brain pericyte proliferation, but the precise molecular mechanisms remain incompletely understood.(Zi *et al*., 2024) Our work identifies Jag2b as a key endothelial ligand linking EC-intrinsic Notch activity to brain pericyte expansion, thereby refining this pathway. How hemodynamics upregulate endothelial Notch signaling, and through which downstream effectors pericyte-intrinsic Notch promotes proliferation, remain to be elucidated. We also found that EC-derived Jag2b contributes to pericyte fate specification. Notably, pericerebral vessels are already perfused before pericytes appear,(Ando *et al*., 2016; Ando *et al*., 2019) raising the intriguing question of whether blood flow regulates fate specification via a similar mechanism.

The opposing roles of Jag2b and Dll4 suggest that brain pericyte proliferation is controlled by a balance between ligand-dependent pro- and anti-expansive influences, thereby ensuring appropriate vascular coverage while preventing excessive pericyte expansion. Disruption of this balance may contribute to cerebrovascular disease. We also found that Jag2b and Dll4 exert opposing effects on cerebrovascular growth. This observation is reminiscent of the opposing effects of Jagged1 and Dll4 reported during retinal angiogenesis.(Benedito *et al*, 2009; Pitulescu *et al*, 2017) Together, these findings raise the possibility that ligand-specific antagonistic interactions represent a broader feature of vascular Notch regulation. Future studies will be required to determine how these ligands differentially activate Notch signaling in pericytes and coordinate pericyte population expansion.

The ligand-specific regulation identified here may also extend to mammalian cerebrovascular development. Mammalian brain vascular cells express multiple Notch ligands, including JAG1, JAG2, and DLL4, whereas NOTCH3 is prominently expressed in vascular mural cells.(High *et al*, 2008; Liu *et al*, 2009; Liu *et al*., 2010; Suzuki *et al*, 2024) Genetic studies have further demonstrated that endothelial Jag1 is required for vascular smooth muscle development and that endothelial JAG1 promotes mural-cell differentiation and NOTCH3 expression, supporting a conserved role for endothelial-to-mural-cell Notch communication.(High *et al*., 2008; Liu *et al*., 2009; Liu *et al*., 2010) These observations raise the possibility that distinct endothelial Jagged and Delta-like ligands may similarly provide differential regulatory inputs to mammalian brain mural cells. However, zebrafish Jag2b cannot be directly equated with mammalian JAG1 or JAG2, and whether Jagged and DLL4 ligands exert opposing effects on mammalian brain pericyte development remains to be determined. Future studies using endothelial-specific manipulation of JAG1, JAG2, and DLL4 in mammalian cerebrovascular models will therefore be important for determining whether the ligand-specific regulatory logic identified here is evolutionarily conserved. Pericyte loss is associated with multiple cerebrovascular and neurodegenerative disorders, raising interest in mechanisms that regulate pericyte maintenance and expansion. Our findings show that altering an endothelial Notch ligand can modify brain pericyte abundance during development and provide a framework for testing whether analogous ligand-specific mechanisms are re-engaged during vascular injury, regeneration, or adult homeostasis. Establishing whether endothelial Jagged/Dll4 signaling can be safely and selectively modulated in mammalian disease models will be necessary before therapeutic implications can be considered.

### Limitations of the study

First, although Jag2b is identified as a key regulator of brain pericyte fate, the specific progenitor populations and Notch receptor(s) involved remain undefined. Second, the severe specification defect in homozygous *jag2b* mutants precluded a direct assessment of Jag2b in later pericyte ingress, and stage-specific genetic manipulation will be required to resolve this function. Third, manipulation of Jag2b, Dll4 and endothelial Notch signaling also alters cerebrovascular development, and indirect endothelial contributions to some pericyte phenotypes cannot be completely excluded. Fourth, the dominant *NICD* and *dnMAML* transgenic approaches produce mosaic and potentially non-physiological pathway perturbations; we therefore interpret these experiments together with complementary genetic, pharmacological and reporter-based evidence. Finally, the mechanisms described here have been defined in zebrafish, and whether analogous ligand-specific regulation operates in mammalian cerebrovascular development remains to be determined.

## Materials and methods

### Zebrafish husbandry and zebrafish lines

Zebrafish (*Danio rerio*) were raised and maintained following the standard procedure in ZFIN in accordance with the Guangdong Provincial Regulations on the Administration of Laboratory Animals, under approval from the institutional animal care and use committee of Guangdong Medical University. The adult zebrafish were maintained in an automated fish housing system (Xinjing Biotech, Shanghai, China) at a temperature of 28°C, with a 14-hour light/10-hour dark cycle. The embryos were kept in an incubator under the same temperature and light/dark cycle. Methylene blue was applied before 2 dpf.

Zebrafish between 2 and 4.5 dpf were used for imaging. As the sex of zebrafish at these stages cannot be determined, there was no sex difference among the experimental animals. Male fish were used for imaging of adult brains.

The following lines were used: *Tg(kdrl:HsHRAS-mCherry)^s896^* abbreviated as *Tg(kdrl:ras-mCherry)*,(Chi *et al*, 2008) *Tg(kdr:EGFP)^s843^* abbreviated as *Tg(kdrl:EGFP)*,(Jin *et al*, 2005) *Ki(pdgfrb-P2A-GAL4-VP16)^ion33d^* abbreviated as *Ki(pdgfrb:GAL4-VP16)*,(Zi *et al*., 2024) *Tg(4xnrUAS:GFP)^c369^* abbreviated as *Tg(4xnrUAS:GFP)*,(Akitake *et al*, 2011) *Tg(fli1ep:dsredex)^um13^* abbreviated as *Tg(fli1a:DsRedEx*),(Lorent *et al*, 2010) *Ki(jag2b-Flag-P2A-EGFP)* abbreviated as *Ki(jag2b-EGFP)*,(Jin *et al*., 2022) *Tg(EPV.TP1-Mmu.Hbb:d2GFP)^mw43^* abbreviated as *Tg(Tp1:d2GFP)*,(Han *et al*, 2016) *Tg(kdrl:tetoff-Hsa.H2B-mScarlet-P2A- dnHsa.MAML1)* abbreviated as *Tg(kdrl:tetoff-H2B-mScarlet-P2A-dnMAML)*, *Tg(kdrl:tetoff-Hsa.H2B-mScarlet-P2A-notch1a-ICD)* abbreviated as *Tg(kdrl:tetoff- H2B-mScarlet-P2A-NICD)*, *Tg(kdrl:Hsa.H2B-mScarlet-P2A-jag2b)* abbreviated as *Tg(kdrl: H2B-mScarlet-P2A-jag2b)*, *Tg(kdrl:Hsa.H2B-mScarlet-P2A-dll4)* abbreviated as *Tg(kdrl: H2B-mScarlet-P2A-dll4)*, *jag2b^+4/+4^* abbreviated as *jag2b^-/-^*, *dll4^+1/+1^* abbreviated as *dll4^-/-^*.

### Processing of scRNA-seq data

Raw sequencing reads were generated in our previous studies. (Li *et al*., 2026; Sun *et al*., 2026) Data from the two experiments were integrated for joint analysis. Quality control was performed using the Seurat package (v5.4.0) in R (v4.4.0). The filtered data were then subjected to unsupervised clustering and visualized using t-distributed stochastic neighbor embedding (t-SNE). To analyze intercellular communication, we used the CellChat package (v1.6.1).(Jin *et al*, 2021) A CellChat object was constructed from the normalized expression matrix, followed by identification of overexpressed genes and ligand–receptor interactions. Potential signaling interactions were inferred, and the cell-cell communication network was summarized using the aggregateNet function to reveal signaling interactions among different cell populations. Further analyses were performed using Loupe Browser (v9.0).

### Genomic DNA purification and genotyping

Genomic DNA was extracted from frozen larvae or tail samples by incubating the tissue in 25 mM NaOH at 95 °C for 30 min, using 25 μL solution per larva or tail sample. An equal volume of Tris-HCl (pH 8.8) was then added to neutralize the NaOH. PCR amplification was performed using 2× Taq PCR Mix (Sangon) with appropriate primers. Genotyping results were analyzed by agarose gel electrophoresis and confirmed by sequencing.

### Generation of knockout lines

The *jag2b^+4/+4^* and *dll4^+1/+1^* lines used in this work were generated by CRISPR/Cas9 technology as described in a previous report. The candidate sgRNAs targeting the exon of *jag2b* or *dll4* were selected by using the online tool CRISPOR (website: http://crispor.tefor.net/crispor.py). The selected sgRNAs were synthesized by Tsingke Biotech (Beijing, China), with modification of 2’-O-methyl and phosphorothioate at the first three 5’ and 3’ terminal RNA residues. To generate the knockout lines, 600 pg Cas9 nuclease (NEB, M0386S) and 125 pg individual candidate sgRNA were mixed and injected into the one-cell-stage embryos of the *Ki(pdgfrb:GAL4- VP16);Tg(4×nrUAS:GFP);Tg(kdrl:ras-mCherry)* line, Next, the mutation rate was examined by sequencing the PCR products containing the mutation area. Injected larvae with high mutation rates were raised to adulthood. Founders with germline transmission of the mutation were identified through outcrossing. F1 larvae carrying heterozygous mutations were raised to establish a stable line.

### CRISPR-mediated gene disruption

For CRISPR-mediated gene disruption in F0 larvae, Cas9 nuclease and gene-specific sgRNA were co-injected into one-cell-stage embryos. *piezo1* crispants were generated using *piezo1* sgRNA, whereas *jag2b* crispants were generated by injecting *jag2b* sgRNA into embryos from *dll4* mutant crosses. Control embryos were injected with Cas9 alone. The sgRNA target sequences are listed in Supplementary file 2. To generate the crispants, 600 pg Cas9 nuclease (NEB, M0386S) and 125 pg individual candidate sgRNA were mixed and injected into the one-cell-stage embryos.

### Generation of *jag2b* and *dll4* cDNAs

Total RNA was extracted from adult zebrafish brain tissue using TRIzol reagent, and purified by using chloroform and isopropyl alcohol according to the manufacturer’s instructions. The mRNA in the total RNA sample was reverse transcribed into cDNA using the 1st Strand cDNA Synthesis Kit (Yeasen). The coding fragment of *jag2b* was amplified by PCR using the primer pair *jag2b*-cDNA F/*jag2b*-cDNA R, and the *dll4* cDNA fragment was amplified using *dll4*-cDNA F/*dll4*-cDNA R. The resulting PCR products were subsequently used for downstream cloning.

### Generation of transgenic lines

The *Tg(kdrl:tetoff-H2B-mScarlet-P2A-NICD)* line, *Tg(kdrl:tetoff-H2B-mScarlet-P2A- dnMAML)* line, *Tg(kdrl:H2B-mScarlet-P2A-jag2b)* line, and *Tg(kdrl:H2B-mScarlet- P2A-dll4)* line were generated by using Tol2-mediated transgenesis.(Bussmann & Schulte-Merker, 2011) *Tg(kdrl:tetoff-H2B-mScarlet-P2A-NICD)* line and *Tg(kdrl:tetoff-H2B-mScarlet-P2A-dnMAML)* line were generated using the plasmids that we reported previously.(Zi *et al*., 2024) *Tol2-kdrl-H2B-mScarlet-P2A-jag2b* and *Tol2-kdrl-H2B-mScarlet-P2A-dll4* plasmids were generated by replacing *jag1b* cDNA with *jag2b* and *dll4* cDNAs in the *Tol2-kdrl-H2B-mScarlet-P2A-jag1b* plasmid. To generate the transgenic lines, 20 pg plasmid and 12.5 pg modified *Tol2* mRNA were mixed and injected into one-cell-stage embryos,(Wang *et al*, 2024) and the larvae with fluorescence expression were raised to adulthood. Adult fish were outcrossed to screen the founder with germline transmission.

### In vivo imaging

*In vivo* fluorescence imaging of zebrafish larvae was performed at room temperature using an FV3000 confocal microscope (Olympus, Japan). Live larvae were anesthetized with 0.02% MS-222 and immobilized in 1.5% low-melting-point agarose in 35-mm glass-bottom dishes (MatTek). Confocal z-stacks were acquired at a z-step of 5 μm with an image resolution of 800 × 800 pixels. For time-lapse imaging, z-stacks spanning the entire imaged cerebrovascular region were acquired from 3.0 to 3.5 dpf at 2-h intervals for a total duration of 12 h for tracking pericyte proliferation. For time-lapse analysis of pericyte specification, z-stacks spanning the entire imaged pericerebral vascular region were acquired from 2.0 to 2.5 dpf. Each z-stack covered approximately 150 μm along the dorsoventral axis with a z-step of 5 μm. Identical imaging settings were used across experimental groups. Images were analyzed using ImageJ.

### Drug application

Yoda1 stock solution was prepared at 10 mM in DMSO and used at a final concentration of 1 μM to activate Piezo1. GsMTx4 stock solution was prepared at 1 mM in ddH₂O and used at a final concentration of 0.1 μM to inhibit Piezo1 activity. DAPT stock solution was prepared at 50 mM in DMSO and used at a final concentration of 50 μM to inhibit Notch signaling. AG1295 stock solution was prepared at 25 mM in DMSO and used at a final concentration of 25 μM to inhibit PDGFRb signaling. Depending on the experimental endpoint, drugs were applied during different developmental windows. For analyses of brain pericyte number, brain vessel segment number, endothelial tip cell number, and pericyte coverage of brain vessels, drugs were applied from 3.5 to 4.5 dpf. For time-lapse analyses of brain ingress of pericerebral pericytes and brain pericyte proliferation, drugs were applied from 3.0 to 3.5 dpf. For analyses of pericyte specification within pericerebral reservoirs, drugs were applied from 1.5 to 2.5 dpf.

### Imaging of adult brains

For imaging of adult zebrafish brains, we employed the CUBIC brain clearing method as previously reported.(Susaki *et al*, 2015; Zi *et al*., 2024) Briefly, brains were dissected and fixed in 4% paraformaldehyde (PFA) at 4 °C for 12 h, followed by washing in cold phosphate-buffered saline (PBS) at 4 °C for an additional 12 h. Samples were then incubated in CUBIC-R1 solution (25% urea, 25% EDTP, and 15% Triton X-100) at 37 °C for 24 h. Cleared brains were embedded in fresh CUBIC-R1 in 35-mm glass- bottom dishes and imaged using an FV3000 confocal microscope (Olympus, Japan). Imaging data were analyzed using ImageJ.

### Cell dissociation and flow cytometry

Approximately 100 zebrafish larvae were used for each biological sample. Larvae were anesthetized with 0.02% tricaine and transferred to pre-chilled DMEM/F12 medium. Brains were rapidly dissected on ice using fine forceps in batches of 5–10 larvae to minimize tissue deterioration. Dissected brains were enzymatically dissociated in DMEM/F12 containing 28 U/mL papain, 0.02 mg/mL DNase, and 0.48 mg/mL L- cysteine, and subsequently washed with DPBS containing 5 mM HEPES, 5% FBS, and 0.29% D-glucose, following a previously described protocol.(Zhang *et al*, 2021) Single-cell suspensions were obtained by gentle trituration followed by filtration. Target cells were isolated by fluorescence-activated cell sorting (FACS) using a Sony MA900 cell sorter (Sony Biotechnology). For analysis of *tyms* and *pcna* expression, EGFP^+^ brain pericytes were isolated from *Tg(kdrl:H2B-mScarlet-P2A- jag2b);Ki(pdgfrb:GAL4-VP16);Tg(4×nrUAS:GFP)* larvae and from *Ki(pdgfrb:GAL4- VP16);Tg(4×nrUAS:GFP)* control larvae at 4.0 dpf. For analysis of endogenous *jag2b* expression, mScarlet^+^EGFP*+* brain ECs were isolated from *Tg(kdrl:tetoff-H2B- mScarlet-P2A-NICD);Tg(kdrl:EGFP)* larvae, whereas EGFP^+^ brain ECs were isolated in parallel from *Tg(kdrl:EGFP)* control larvae at 6.0 dpf. Approximately 300 sorted cells were collected per biological sample for reverse transcription and pre- amplification.

### Reverse transcription and quantitative real-time PCR (qPCR)

cDNA synthesis and pre-amplification were performed directly from the sorted cells using the SuperScript IV Single Cell/Low-Input cDNA PreAmp Kit (Thermo Fisher Scientific) according to the manufacturer’s instructions. The resulting pre-amplified cDNA was subjected to quantitative real-time PCR (qPCR) using TB Green Fast qPCR Mix (Takara Bio). Endogenous *jag2b* expression was quantified in FACS-isolated brain ECs from EC-*NICD* and control larvae, whereas tyms and pcna expression was quantified in FACS-isolated brain pericytes from EC-*jag2b*-overexpressing and control larvae. Relative gene expression was calculated using the comparative Ct method after normalization to the reference gene *eef1a1l1*. Three independent biological replicates were analyzed for each comparison. Statistical significance between two groups was assessed using an unpaired two-tailed Student’s t-test applied to the ΔCt values.

### Quantification of brain pericytes, brain vessels and pericyte coverage of brain vessels

To quantify brain pericyte population, the brain pericyte number and the pericyte coverage of brain vessels in the midbrain were measured in this study. The Cell Counter function of ImageJ was used to manually enumerate the brain pericyte number and the segment number of the brain vessels in *Ki(pdgfrb:GAL4- VP16);Tg(4×nrUAS:GFP);Tg(kdrl:ras-mCherry)* larvae. In this study, only the pericytes that cover the brain vessels and their cell bodies completely disengaged from pericerebral vessels could be defined as brain pericytes. The density of pericytes on brain vessels was quantified by calculating the pericyte coverage of brain vessels, following the approach previously described,^37^ which was calculated as the brain pericyte number divided by the number of brain vessel segments. All analyses were performed blinded to genotype and treatment.

### Quantification of pericyte proliferation and brain ingress of pericerebral pericytes

Time-lapse imaging from 3.0-3.5 dpf was used to quantify brain pericyte proliferation as previously described.(Zi *et al*., 2024) In this work, analyses were restricted to pericytes in the midbrain. Brain pericytes at the start and end of imaging were counted in ImageJ, and the origin of newly appearing pericytes was traced. Migration of a pericyte from a pericerebral vessel into the brain was defined as an ingress event, whereas division of a brain pericyte producing two daughter cells was defined as a proliferation event. Proliferation frequency was calculated as the total number of proliferation events within 12 h divided by the initial number of brain pericytes. Specifically, a pericyte initially located on a pericerebral vessel at the start of imaging that subsequently migrated along brain vessels into the brain was classified as an ingress event. All analyses were performed blinded to genotype.

### Statistics

Zebrafish larvae were randomly assigned to experimental groups, and image quantification was performed blinded to genotype and treatment. The significance of the difference between two groups was calculated using an unpaired two-tailed Student’s t-test, which was performed using the GraphPad Prism v9.0 software. The results were represented as mean ± SEM in all figures, and the level of significance was indicated as follows: **** indicates p < 0.0001, *** indicates p < 0.001, ** indicates p < 0.01, * indicates p < 0.05, and n.s. indicates p > 0.05.

## Acknowledgements

We thank Dr. Jiulin Du for providing the *Ki(pdgfrb:GAL4-VP16);Tg(4×nrUAS:GFP)* and *Tg(Tp1:d2GFP)* lines. We thank Dr. Jie He for providing the *Ki(jag2b-EGFP)* line. We thank Dr. Jia Li for help with the generation of the *Tol2-kdrl-tetoff-NICD* and *Tol2- kdrl-tetoff-dnMAML* plasmids. This work was supported by the National Natural Science Foundation of China (31701265) and the National Key R&D Program of China (2024YFA1802200) and Guangdong Provincial Project for Teaching Quality and Teaching Reform in Undergraduate Institutions (Yue Jiao Gao Han [2024] No.9, No.30), China.

## Additional information

### Funding

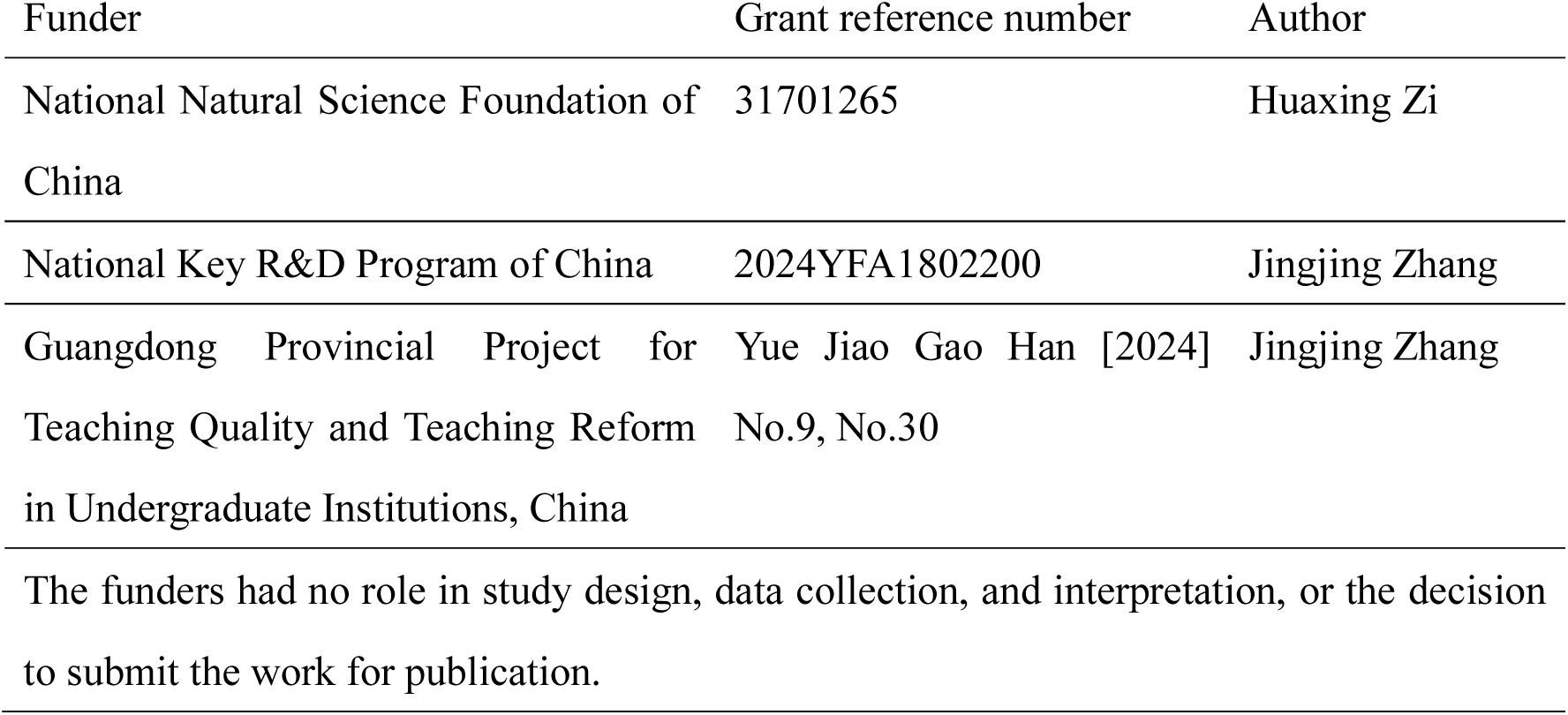

### Author contributions

Jingjing Zhang, Conceptualization, Supervision, Writing – review & editing. Huaxing Zi, Conceptualization, Methodology, Investigation, Writing – original draft, Writing – review & editing. Chi Dang, Methodology, Investigation; Xiaoyu Li, Formal analysis; Weiyi Sima, Investigation, Resources; Guozhu Ning, Investigation; Qiang Wang, Resources; Li Sun, Resources.

### Ethics

Zebrafish were raised and maintained according to standard ZFIN procedures and in accordance with the Guangdong Provincial Regulations on the Administration of Laboratory Animals. All animal procedures were approved by the Institutional Animal Care and Use Committee of Guangdong Medical University.

## Additional files

### Supplementary files

• Figure supplements and corresponding legends for Figures 1–6.

• Legends for supplement videos.

• Supplementary file 1. Key resources table

• Supplementary file 2. Oligonucleotide sequences.

### Data availability

The previously published scRNA-seq datasets analyzed in this study are publicly available in the Gene Expression Omnibus (GEO) under accession GSE243907 and in the Genome Sequence Archive (GSA) under accession CRA030225. All data generated or analyzed during this study are included in the manuscript and its supporting files. This study did not generate new sequencing datasets or original code.

